# Network structural features affect stability of Calvin Bassham Benson (CBB) pathway in plants

**DOI:** 10.1101/034637

**Authors:** Matthew K Theisen

## Abstract

The stability of the Calvin Bassham Benson (CBB) cycle remains an area of active computational research. Our understanding of biology and the prospect for bioengineered plants with higher productivity may both be impacted by a greater understanding of this area. Here we use the ensemble modelling robustness analysis (EMRA) framework to show that the action of the phosphate/G3P antiporter is much more significant for maintenance of stability than a recently proposed G6P shunt. Additionally, we interpret recent results suggesting that overexpression of RuBiSCO does not improve growth rate of plants but overexpression of SBPase does. Our simulations reproduce this result, but only in models which do not include the G6P shunt. Taken together, these results may suggest a situational role for the G6P shunt, possibly in dynamic situations under starvation or other stress conditions.

## Introduction

The Calvin Bassham Benson cycle (CBB) is responsible for CO_2_ fixation by plants, including the C3 & C4 variants, of which the C4 is an adaptation which allows for plants in high temperature or low water environments (1–4). Plants have advanced regulatory systems which allow them to successfully grow and thrive in an unpredictable and changing world (5–9). For example, sugars generated from CO_2_ during daylight are stored as starch in photosynthetic and non-photosynthetic chloroplasts. Nighttime consumption of starch is tuned to leave only a small amount remaining by morning—and this consumption rate is dynamically tuned to adjust for changing day length(10–12).

Among canonical metabolic pathways, the CBB pathway is highly branched and complex, much more so than simple linear pathways like glycolysis or simple loops like the TCA cycle. This is in some ways the result of the chemical difficulty of aerobic CO_2_ fixation (13,14) which seems to require a carbon reshuffling step to regenerate a suitable starting substrate like ribulose-1,5-bisphosphate (15). The complexity of the pathway, in which there is not a linear pathway from substrate to product, results in instability if intermediates are depleted. For example, if the sugar phosphates in a chloroplast become depleted, the pathway is not able to continue since some starting substrate (RuBP) is required to continue CO_2_ fixation (16).

There are two main mechanisms of transport of sugars across the chloroplast membrane. First, there is the G3P/phosphate antiporter (17). This transporter moves a G3P from the CBB pathway in the chloroplast to the cytosol, where it is used for various cellular functions. In return a phosphate molecule is transported into the chloroplast, effectively keeping the total number of phosphates in the chloroplast constant. Second, there are glucose (putative) (18) and maltose transporters (19), of which the maltose transporter is known to be essential for starch breakdown. G3P is a CBB intermediate and is directly interconvertible with other sugar phosphates, so a depletion of G3P would be problematic for CBB. However, glucose and maltose are more removed from the CBB pathway itself and are possibly only produced as starch breakdown products (12).

The direct regulation of plastidic enzymes involved in photosynthesis is accomplished by redox-mediated proteins called thioredoxins (20). In light conditions, the NADPH/NADP^+^ ratio is higher because the photosystems which generate NADPH from light are active. As a result, the disulfide bonds in thioredoxins and other regulated proteins are broken, mediating enzyme activity. In Arabidopsis thaliana several enzymes are known to be redox regulated in this manner (21).

Some enzymes of the CBB cycle are activated in a reducing (light) environment by the breaking of their disulfide bonds. In the dark, these enzymes are attenuated in the oxidizing environment. Of the 12 enzymes of the canonical CBB cycle, 4 are known to be redox regulated in the ferredoxin/thioredoxin system (21). First, GAPDH converts 1,3-bisphosphoglycerate to glyceraldehyde-3-phosphate using reducing power from NADPH. GAPDH is reversible, although in dark conditions scarce NADPH is required for other critical cellular functions.

In addition to GAPDH, enzymes which catalyze the cleavage of high-energy phosphate bonds are also thioredoxin-regulated, presumably to reduce thermodynamic losses in dark conditions. Phosphoribulokinase (Prk) catalyzes the cleavage of ATP to ADP coupled with the conversion of ribulose-5-phosphate to ribulose-1,5-bisphosphate. Prk and GAPDH are inactivated in the non-enzymatic oligomerization of with CP12 in oxidizing conditions, which is reversed by NADPH (22). Sedoheptulose-1,7-bisphosphatase catalyzes the irreversible loss of phosphate from the sedoheptulos-1,7-bisphosphate to result in sedoheptulose-7-phosphate. Fructose-1,6-bisphosphatase catalyzes an analogous reaction and loss of phosphate to result in F6P. These enzymes are all regulated to lose function in dark conditions when NADPH is low and CO2 fixation cannot continue (21).

The Calvin cycle has many enzymes in common with the pentose phosphate pathway, except that it functions in the reverse direction, leading to the distinction between the traditional or oxidative pentose phosphate pathway (oPPP) and the CBB-synonymous reductive pentose phosphate pathway (rPPP) (23). Distribution of oPPP and rPPP enzymes within plant cellular compartments (plastid vs. cytosol) is an area of research (24), but in *Arabidopsis*, it is recognized that the first three steps of the oPPP (glucose-6-phosphate dehydrogenase, gluconolactonase and 6-phosphogluconate dehydrogenase) are localized to both the plastids and the cytosol (25). In addition to the CBB enzymes above, the plastidic enzymes of the oPPP, particularly G6PDH, are subject to redox-based regulation (26).

G6PDH is most active in oxidizing conditions which prevail in night darkness. The oPPP provides NADPH for critical cell functions when light is unavailable. Activity is highly attenuated by the presence of light. This is rationalized to be for the prevention of thermodynamic losses due to a futile cycle (27). However, interestingly, the attenuation of G6PDH in reducing conditions is far from complete and varies widely by species. In the investigation of three different plastidic G6PDHs, activity is attenuated to anywhere from 10-30% of maximum in reducing conditions (26,28,29). It has recently been suggested that flux through G6PDH and the next two oPPP enzymes (generating Ru5P) may stabilize the CBB pathway (30). This opens the door for investigation into possible competitive benefits of a futile cycle which in terms of thermodynamics, is a clear loss.

Some previous efforts have attempted to address stability in the CBB pathway, but these have had shortcomings such as not considering phosphate (31) which our work suggests has a critical role in stability, or considering only a single set of parameter values (32), which doesn’t reflect the range of stochastic and environmental variability encountered in biological reality. Other works focused on the well-documented oscillations of the CBB pathway (33), without considering general propensity towards stability (nonsingular Jacobian), or instability (singular Jacobian). In this work, we consider the present evidence that multiple structural features of CBB in plants and *Arabidopsis thaliana* in particular stabilize the pathway, independent of their effect on oscillatory behavior. In particular, we investigate the role of the G3P/phosphate translocator, the oxidative pentose phosphate pathway, as well as covalent modification of triose phosphate isomerase (34). We use ensemble modeling robustness analysis, a method which investigates the stability of metabolic pathways using network information such as reference flux, network stoichiometry, reaction reversibility and substrate-level regulations (35). We also consider the potential applications toward biotechnological work attempting to increase the productivity and growth rates of plants.

## Chloroplast model

First, a consensus model of chloroplast metabolism flow in light conditions was developed (Fig. 1). Steps of the CBB cycle, starch synthesis and starch degradation were included. Additionally, G3P transport from chloroplast to cytosol is also included. NADPH generation by the light reactions and ATP generation through respiration were included as single reactions in the model.

**Figure 1.**
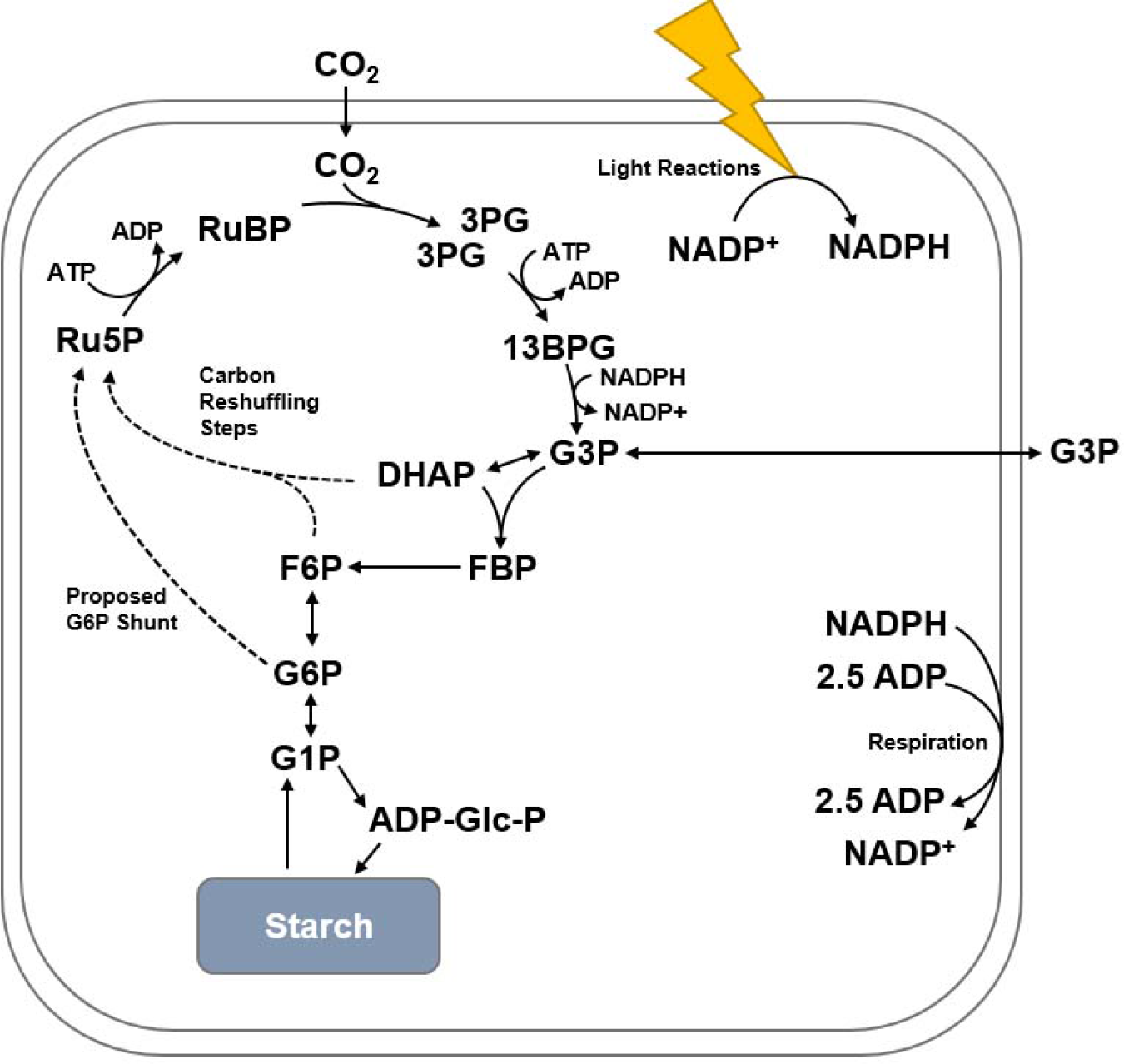
The overall model of chloroplast flux used in this paper. Reactions in the carbon shuffling steps and G6P shunt are modelled individually but are shown in a simplified format. Light reactions and respiration were modeled as single reactions for simplicity. NADPH/NADP+ & ATP/ADP cofactors included in all simulations, free phosphate held constant in some simulations as noted. CO_2_ was held constant in all simulations.

Fluxes were set by linear programming to determine a reference steady state. Carbon was assumed to be split 50:50 between G3P and starch synthesis. Starch degradation was assumed to be at 2/3 the rate of starch synthesis. Starch degradation is represented as non-negligible in the model, since starch degradation rate was found to be almost unchanged by light in spinach leaves (36). G3P export and import were modeled as parallel reactions in dynamic equilibrium, additionally at a 2:3 ratio for parsimony. Beyond these specifications, the system has no degrees of freedom so flux rate was completely determined.

Reactions were modeled kinetically using realistic rate laws which take into account number of substrates and products, and the reversibility of the reaction using modular reaction rates according to the method of Liebermeister (37). Substrate-level regulations were added to the model as described in the Methods section. Parameter values were sampled constrained to reference fluxes and stability at the reference steady state was insured. A suitable number of parameter value sets (n = 300) were generated and tested. Enzyme levels were perturbed by using the parameter continuation method, where the system is perturbed, constrained to a fixed point, until the Jacobian becomes singular, or a metabolite concentration becomes negative. The fraction of parameter sets, or ‘models’ which become unstable at each level of integration, is plotted. Further details about the model are available in the methods section.

## Results

### G3P/phosphate translocator almost completely stabilizes CBB

The G3P/phosphate translocator and phosphate in general is known to have an important role in the action of the CBB pathway (Fig. 2A) (38). However, to date, this role has not been thoroughly tested by simulation efforts. Here we test the idea of the phosphate antiporter as a CBB pathway stabilizer by doing EMRA simulations of the CBB enzymes with and without holding phosphate constant. Allowing plastidic phosphate to vary freely as a metabolite in the simulation is a proxy for the effects of the antiporter, since if phosphate was transported independently from the cytosol, there would be effectively no steady state requirement for phosphate—any deviation would simply be made up by transport to or from the cytosol. The inclusion of phosphate fixes the steady state requirement to the one-to-one antiport of G3P and inorganic phosphate.

**Figure 2.**
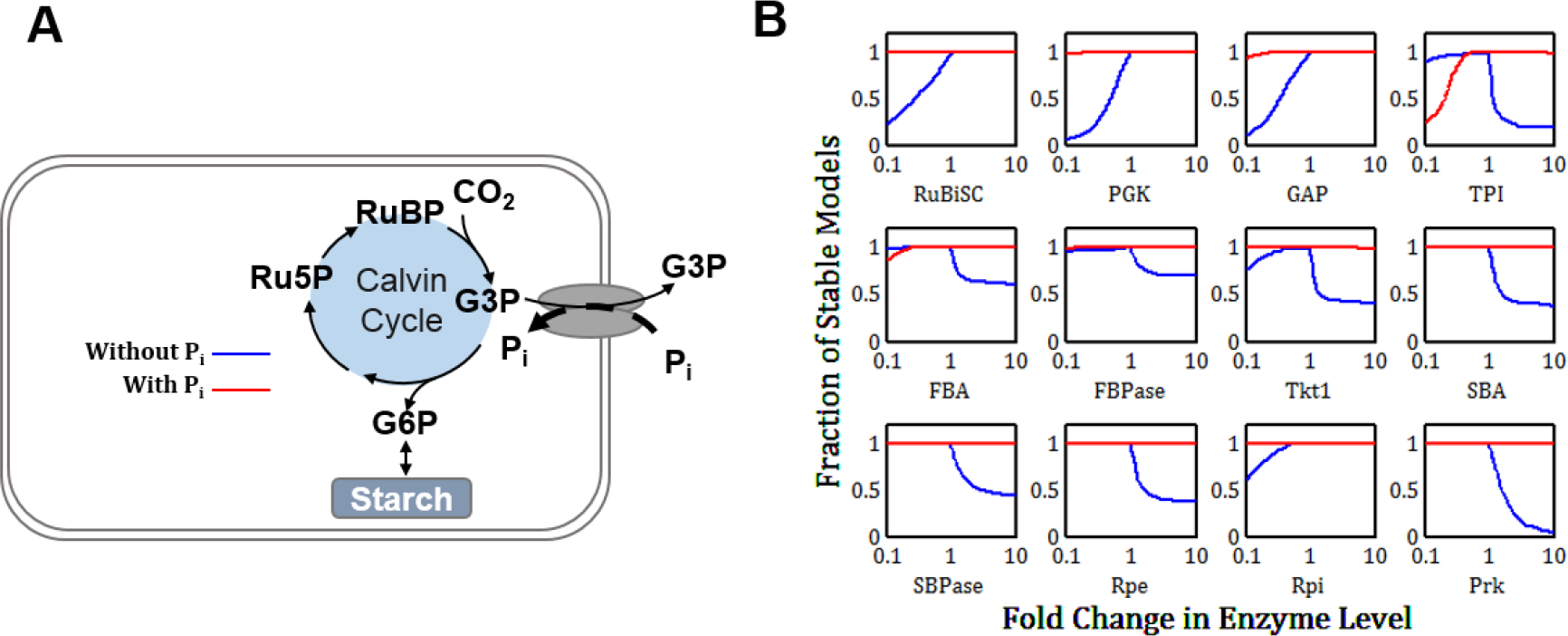
Comparison of stability with and without holding phosphate constant. **A)** Schematic showing the flow of phosphate through the phosphate/G3P antiporter in relation to the CBB pathway. **B)** EMRA stability profile for the enzymes of the CBB pathway upon perturbation of 10x and 0.1x. Both Tkt reactions were perturbed simultaneously (n = 300). Including the effects of the G3P/phosphate antiporter (red line) significantly stabilizes the pathway.

Without P_i_, the model was found to have noticeable instability in essentially all of the CBB enzymes, and, seven of the 12 CBB enzymes were noted to have instability upon increase. Inclusion of phosphate as a metabolite was shown to almost completely eliminate instability with one notable exception. With phosphate, triose phosphate isomerase was noticeably unstable to decrease. Without phosphate, that enzyme was unstable to increase—the inclusion of phosphate reversed the tendency towards instability (Fig. 2B, red lines).

### Glutathionylation increases stability of TPI in phosphate condition

The one enzyme of the CBB pathway which was unstable after the inclusion of phosphate was triose-phosphate isomerase. There seems to be further experimental confirmation of the importance of sufficient TPI activity. A plastidic TPI mutant with reduced activity was installed in *Arabidopsis* and the resulting plants were found to grow at a highly stunted rate (39). Interestingly, if grown in the dark with nutrients provided (heterotrophic growth), there was no growth deficiency, indicating that the plastidic TPI is important for autotrophic (light) metabolism, but not critical for heterotrophic (dark) metabolism.

There are multiple possible methods for accommodating this loss of stability. First, triose-phosphate isomerase is a highly active, reversible enzyme with no stability penalty indicated (Fig. 2B) for high activity, so it’s possible TPI is operating mostly or exclusively in the high activity domain, where stability is not an issue. Another possibility is that TPI instability is partly rescued by the effect of glutathionylation. A recent analysis showed the first evidence of glutathionylation of plant enzymes. The authors found that a cytosolic TPI from Arabidopsis thaliana was inactivated in the presence of oxidized glutathione (GSSG) but reactivated in the presence of reduced glutathione (34). Since GSFI is regenerated by the reducing power of NADPH, this regulatory network can be represented as NADPH activation of TPI combined with NADP^+^ inactivation (Fig. 3A).

**Figure 3.**
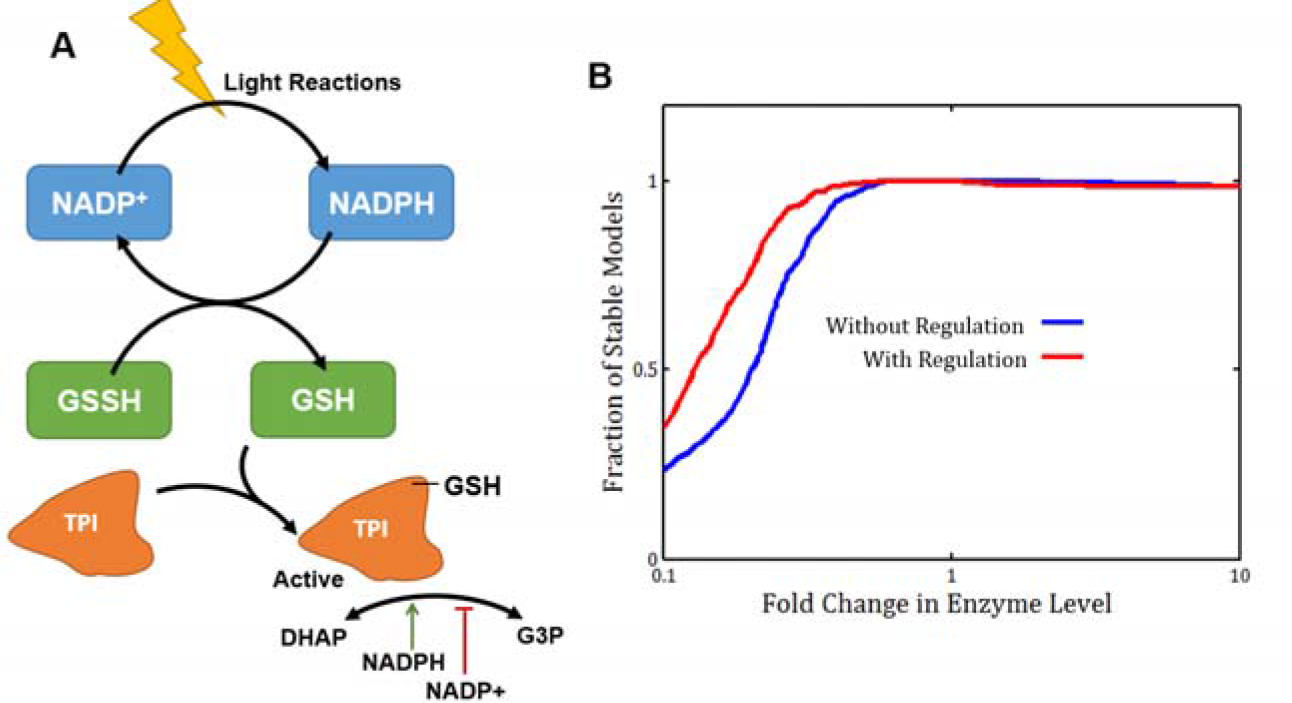
Possible regulatory mechanism for plastidic TPI. **A)** Possible schematic for TPI activation by glutathionylation. This is represented in the model by NADPH activation and NADP^+^ repression. **B)** Stability profile for the TPI enzyme in the ‘with phosphate’ model showing the effect of NAPDH regulation on TPI.

Interestingly, the stability of the TPI with NADPH/NADP+ regulation improves noticeably (Fig. 3B). Although there is no direct evidence if glutathionylation of plastidic TPI (pdTPI) in Arabidopsis, the protein sequences show 62% sequence identity and have similar numbers of methionine residues (2 & 3), (UniProt entries Q9SKP6 & P48491 (40), aligned by BLASTP 2.3.0+ (41,42)). Regulation of plastidic TPI may be an interesting area of future research.

### Glucose-6 phosphate shunt affects stability of no phosphate condition

A perhaps paradoxical aspect of the plastidic glucose 6 phosphate dehydrogenase enzyme is that it retains some activity after deactivation, which seems to be thermodynamically unfavorable, since carbon decarboxylated by the oxidative pentose phosphate pathway has to be re-fixed by RuBiSCO, including the use of 3 ATP per carbon fixed. It has recently been proposed that this is a feature of chloroplastic metabolism which may stabilize the CBB pathway itself (30). This was discussed in great detail but so far has seen no mathematical justification. Looking at the CBB with stabilization by the phosphate translocator, there is little stability improvement to be made. In stress conditions, however, such as phosphate limitation, plant metabolism is known to change radically (43–45), including changing expression of plastidic transporters (46). This could potentially alter the stabilizing, protective effects of the phosphate/G3P antiporter, which can be modelled (as before) by the removal of phosphate as a metabolite. In such cases, other structural features would be required to provide stability.

The so-called glucose-6 phosphate shunt (Fig. 4A) has been proposed to provide stability to the CBB. To test the effects of the proposed glucose-6 phosphate shunt, simulation of the no-phosphate condition with various levels (0% of RubisCO, 10%, 30%) of flux through the first three enzymes of the oxidative pentose phosphate pathway (G6PDH, GLNase & GLNDH) was undertaken via EMRA. For nearly all enzymes, the 10% & 30% conditions showed stability improvements over the 0% condition for increases in enzyme activity from the reference steady state (Fig. 4B, red & green lines). However, interestingly, several enzymes showed slightly higher instability in the 10% and 30% conditions upon *decrease* in enzyme amount, though higher stability upon increase. One possible explanation is that transketolase, and the aldolases are highly active, reversible enzymes, and thus more likely to operate in the high activity regime than the low activity. Another possible explanation is that the operation of the G6P shunt is situational, and that it is meant to operate in dynamic scenarios to replenish cycle intermediates, rather than to operate continuously to maintain steady state.

**Figure 4.**
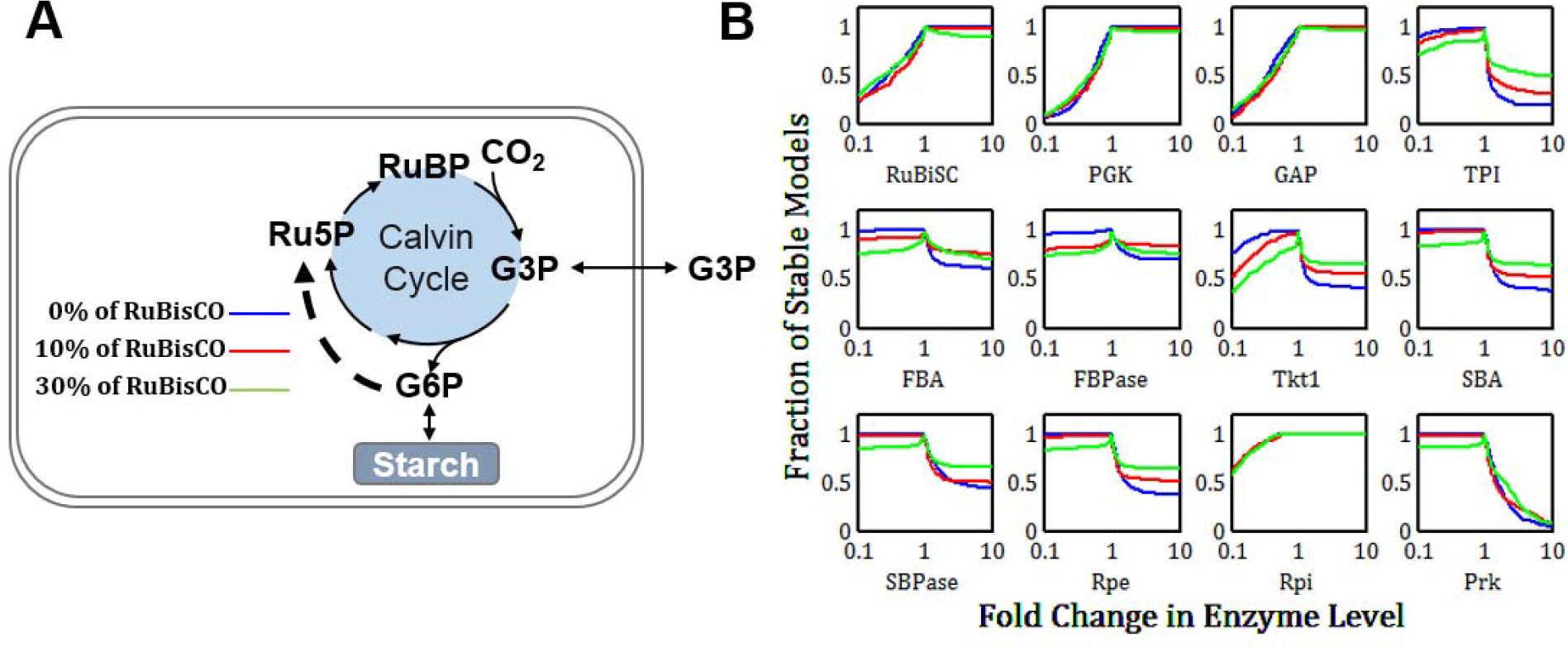
Comparison of stability of various fluxes through the proposed G6P shunt. **A)** Schematic showing the flow of metabolites through the G6P shunt relative to the CBB pathway. **B)** EMRA stability profile for the enzymes of the CBB pathway upon perturbation of 10x and 0.1x. Both Tkt reactions were perturbed simultaneously (n = 300). Including the effects of the G6P shunt (red line & green lines) improves stability of the pathway upon increase of many enzymes, hurts stability upon decrease of many enzymes (Tkt, aldolases, phosphatases particularly).

### Assessing methods for improving plant productivity, SBPase and RuBiSCO overexpression

Use of stability analysis to provide biological insight into the mechanisms of stability in the CBB is one powerful demonstration of its capabilities. However, it doesn’t provide insight into engineering and biotechnological efforts which are aimed at increasing the productivity of plants, particularly relating to growth rate and the CO2-fixing rate of the CBB pathway. Thus, in addition to assessing the effect of genetic changes on stability, we can additionally look at the predicted impact on net carbon fixation rate.

Many efforts to increase growth rate and carbon fixation rate of plants have understandably focused on RuBiSCO. Some projects have focused on methods to modify the amino acid sequence of RuBiSCO (47,48). Others have attempted to overexpress RuBiSCO or, more recently, replace native RuBiSCO with a heterologous enzyme which has higher specific activity (49). These efforts have increased the content and activity of RuBiSCO, but they have not convincingly increased plant productivity (50). However, looking at the CBB pathway as a network problem rather than a problem with a single enzyme opens up many different possibilities. Interestingly, one group reported that overexpression of SBPase increased carbon fixation rate by 6-12% (51).

To investigate consistency of these results with simulation, the average model-predicted net CO2-fixation rate for different genetic changes and flux configurations can be compared. Interestingly, results show that for the 0% G6PDH condition with phosphate, overexpression of SBPase slightly increased CO2-fixation rate, while RuBiSCO overexpression was, counterintuitively, found to decrease RuBiSCO flux. Other targets in the CBB pathway which were also investigated, with Prk showng the largest projected increase on carbon fixation rate (Fig. 5A). For other conditions (no phosphate, with G6PDH flux) (Fig. 5B & C), no improvement was observed for either, except a small improvement for SBPase in the no phosphate model. This suggests that perhaps network effects are more determinative of the response of the CBB pathway than performance of individual enzymes. Additionally, it seems to suggest that in laboratory conditions, the models not including G6P shunt flux are more reflective of biological reality, and thus that the role of the G6P shunt may be situational.

**Figure 5.**
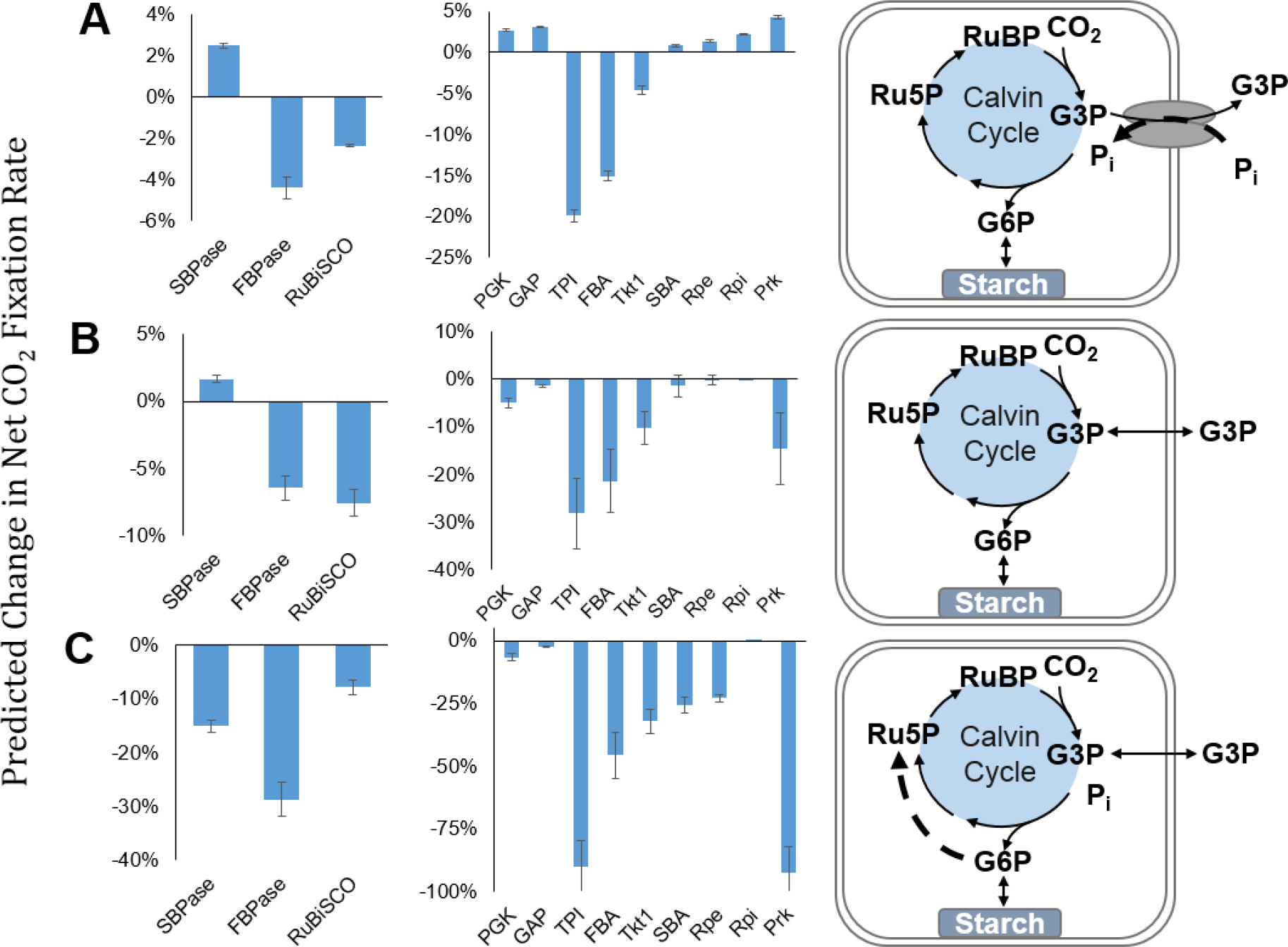
Figure showing predicted effect of 10x increase of various CBB enzymes for three different flux models. SEM shown (n = 300). **A)** Allowing plastidic phosphate to vary freely. **B)** Holding phosphate constant. **C)** With G6P shunt at 30% of RuBiSCO flux.

## Discussion

This analysis reveals the importance of structural features for the stability of the CBB pathway in plants. Stability is an important characteristic of metabolic pathways, since they are subject to stochastic variability in protein expression as well as different environmental conditions which can perturb the system. While oscillations in the CBB are a point of previous research (33), we here present an analysis of the stability of the underlying fixed points involved. So far, stability, and in particular the ensemble modeling robustness analysis framework has been applied to explore the performance relatively simple *in vitro* pathways, but this paper shows how it can also uncover and illuminate biologically significant features and phenomena.

Additionally, this manuscript sheds additional light on some specific details of these mechanisms. For instance, these results indicate that the G3P/phosphate antiporter is more significant for the stability of the CBB than the G6P shunt under normal steady state. However, if the one-to-one link between phosphate- and G3P-transport is broken (as in the no phosphate simulations), the action of the glucose-6-phosphate shunt does change the stability profile of CBB enzymes noticeably. However, the true purpose of the G6P shunt may be to restore steady state in dynamic situations. This sheds light on the apparent paradox of thermodynamic losses in this ‘futile’ cycle. The thermodynamic involved in one turnover of the oPPP would be involve loss of one ATP in the Prk step and two ATP at the Pgk step.

Among heterotrophic organisms using the CBB cycle, there is a remarkable amount of diversity in the arrangement and function of metabolism (52–56). Thus, it is likely that depending on environmental constraints and chance occurrences in evolutionary history, the stabilizing mechanisms used by different species are a combination of those presented here and those yet to be discovered. Thus, this manuscript is not a comprehensive or conclusive look at the mechanisms of stability in the CBB pathway but is an initial, provisional investigation into some possible explanations for the success of the CBB pathway despite its apparently unstable underlying structure. The model presented here is advances on some previously described models (31–33) in important ways. This work provides answers and more questions to pave the way for yet more complete and sophisticated simulation of CBB.

So far, attempts to increase the productivity of plants have mostly focused on individual enzymes, rather than investigating the CBB pathway as a network. Here, we give plausible explanation to results that show SBPase overexpression increases plant growth rate while RuBiSCO overexpression has so far not shown any increase in plan performance. While the methods employed here are not conclusive, they provide new insights which lay out potential targets of future exploration in the biotechnological engineering of plants.

## Methods

The model of chloroplast metabolism, including the CBB, the G3P/phosphate translocator and the was constructed by inspecting the latest literature about plastid metabolism (12). The full stoichiometric matrix, reversibilities and reference flux are shown in Supplementary Table 1. Adjustments were made as necessary (removal of phosphate, adjustment of fluxes to include G6P shunt etc.). Based on stoichiometry and reversibility, realistic Michaelis-Menten style rate laws were assigned. Regulation of PGM, G6PDH were included and TPI was regulated in some simulations. Parameters were obtained by randomly sampling normalized affinity parameters from a uniform distribution (0.1,10) as described previously. V_max_ was then solved for, constraining the rate law to the reference steady state. Simulations of steady state perturbations were carried out using the parameter continuation method described previously (35). Calculations were done in MATLAB and full code with instructions to reproduce all data will be provided.

